# Retinal and Cortical Determinants of Cortical Magnification in Human Albinism

**DOI:** 10.1101/562249

**Authors:** Erica N. Woertz, Melissa A. Wilk, Ethan Duwell, Jedidiah Mathis, Joseph Carroll, Edgar A. DeYoe

**Author notes:** HudsonAlpha Institute for Biotechnology, 601 Genome Way, NW, Huntsville, AL 35806. Corresponding Author: Edgar DeYoe, Department of Radiology, Medical College of Wisconsin, 8701 Watertown Plank Road, Milwaukee, WI 53226.

## Abstract

The human fovea lies at the center of the retina and supports high-acuity vision. In normal visual system development, foveal acuity is correlated with both a high density of cone photoreceptors at this location and a magnified retinotopic representation of the fovea in the visual cortex. Both cone density and the cortical area dedicated to each degree of visual space—the latter known as the cortical magnification function—steadily decline with increasing eccentricity from the fovea. In albinism, peak cone density at the fovea and visual acuity are reduced but appear to be normal in the periphery, thus providing a model to explore the correlation between retinal structure, cortical structure, and behavior. Here, we used adaptive optics scanning light ophthalmoscopy to assess retinal cone density and functional magnetic resonance imaging to measure cortical magnification in primary visual cortex of normal controls and individuals with albinism. We find that retinotopic organization is more varied in albinism than previously appreciated, yet cortical magnification outside the fovea is similar to that in controls. Moreover, cortical magnification in albinism and controls exceeds that which might be predicted based on cone density alone, suggesting that reduced foveal cone density in the albinotic retina may be partially counteracted by central connectivity. Together, these results emphasize that central as well as retinal factors must be included to provide a complete picture of aberrant structure and function in genetic conditions such as albinism.

## INTRODUCTION

The human fovea occupies about 0.02% of the total retinal area, but is responsible for our highest-acuity vision, with approximately 40% of primary visual cortex (V1) dedicated to processing its signals (Hendrickson, 2005). The fovea is characterized by the excavation of inner retina, a lack of retinal vasculature, and a small region absent of rod photoreceptors. The fovea has the highest density of cone photoreceptors, as well as non-convergent connections between these cones and their post-synaptic partners, known as the “midget” system (Curcio and Allen, 1990; Dacey, 1993). This leads to cortical sampling that is up to 160 times greater than that of peripheral cones (Duncan and Boynton, 2003). Cone density declines with increasing eccentricity, with the steepest decline occurring within 1-2 mm of the fovea (Curcio et al., 1990). Likewise, in V1 the amount of cortical space devoted to each degree of visual angle is grossly magnified at the fovea and decreases steadily with increasing eccentricity (Daniel and Whitteridge, 1961; Cowey and Rolls, 1974).

Behaviorally, this relationship between cortical magnification (CM) and eccentricity is directly correlated with visual acuity. This was shown with the earliest empirical measurements of CM in humans using electrodes implanted in the occipital lobe (Cowey and Rolls, 1974). Later studies using functional magnetic resonance imaging (fMRI) showed strikingly similar CM measurements, though there is some minor variability that has been attributed to disparities in stimulus design and methods used to measure cortical distance (Sereno et al., 1995; Engel et al., 1997; Popovic and Sjöstrand, 2001; Duncan and Boynton, 2003; Qiu et al., 2006).

While methodological differences may contribute to this variation in empirical measurement of CM, it is also possible that some variation comes from actual anatomical differences between individuals. In the retina, histologic studies show more than a 3-fold difference in foveal cone density across individuals (Curcio et al., 1990), and the sizes of the optic tract, lateral geniculate nucleus (LGN), and V1 are known to be correlated within individuals (Andrews et al., 1997). Since foveal magnification in the visual cortex is driven in part by the relatively higher cone density found at the fovea, it follows that CM should vary across subjects as a function of cone density (Dougherty et al., 2003). Recent advances in retinal imaging with adaptive optics has made it possible to resolve the cone mosaic in the living human eye, providing the opportunity to assess cone density and CM in the same individual. Such comparisons can determine whether variation in retinal structure correlates with variation in CM.

Extreme examples of individual variation are found certain pathologies. One notable example is albinism, a family of genetic diseases that disrupt melanin synthesis and cellular trafficking, resulting in abnormal development of the visual system (Nettleship, 1909; Creel et al., 1978; O’Donnell et al., 1978). This leads to greater visual system variability among individuals with albinism than among normal individuals, with peak cone density that is (on average) lower than that observed in normal individuals (Wilk et al., 2014). Thus, albinism is an advantageous model to probe structural correlates of variability in CM. Additionally, individuals with albinism are known to experience decreased best-corrected visual acuity compared to normal individuals (Wilson et al., 1988; Summers, 1996). Although they generally have foveal cone densities below the normal range of 100,000-320,000 cones/mm^2^ (Curcio et al., 1990) with minimum foveal cone densities as low as 29,000 cones/mm^2^, their visual acuity and cone density does not appear to be precisely correlated (Wilk et al., 2014). Thus, the physiological source of their acuity deficits remains unclear. Several studies have assessed cortical reorganization in albinism (Morland et al., 2001; Hoffmann et al., 2003; von dem Hagen et al., 2005; von dem Hagen et al., 2007; Neveu et al., 2008; von dem Hagen et al., 2008; Bridge et al., 2014; Ather et al., 2019), but these studies lacked molecular genotyping to confirm the specific subtypes of albinism included in their cohorts. Moreover, to our knowledge no studies in this population have assessed CM and its quantitative relationship to cone density. In light of the above-mentioned relationship between visual acuity and CM in normal individuals, it is of particular interest to determine how CM in albinism compares to that in normal individuals and if it correlates with observed decreases in cone density at different eccentricities. This will provide a bridge between visual system structure and aberrant acuity functions in albinism, as well as insight into the nature of pathological retino-cortical relationships in disease.

Here, we performed high-resolution retinal imaging using adaptive optics scanning light ophthalmoscopy (AOSLO) and retinotopic mapping in the visual cortex using fMRI to examine both cone density and CM in the normal and albinotic visual system. We hypothesized that CM would be reduced in subjects with albinism in proportion to their unique pattern of reduced cone density versus visual field eccentricity. However, our results suggest that cone density is not the sole determinant of CM and its variability. Additional central factors are required to account for the aberrant cortical patterns within this population.

## METHODS

### Subjects

Six subjects with albinism (4 females, 2 males; aged 15-31 years) with minimal nystagmus and five subjects with no prior ocular or cortical pathology (2 females, 3 males; aged 20-25 years) were recruited for this experiment. One subject with albinism was excluded from further analysis due to significant motion artifacts in the fMRI (male, age 15 years); subjects included in the analysis are listed in **Table 1**. Retinal features of these subjects have been previously described (Wilk et al., 2014; Wilk et al., 2017). The study was in accordance with the Declaration of Helsinki and approved by the Institutional Review Board of the Medical College of Wisconsin. All subjects provided written consent after explanation of the nature and possible consequences of the study.

**Table 1.** Subject demographics.

| Condition | Subject | Sex | Age | Eye | Axial Length | Foveal |
| --- | --- | --- | --- | --- | --- | --- |
|  |  |  |  |  | (mm) | Cone Density<br>(cones/mm<sup>2</sup>) |
| Normal | JC_0200 | M | 25 | OD | 24.72 | 128,560* |
|  | JC_0677 | F | 24 | OD | 24.03 | 165,080* |
|  | JC_0769 | F | 21 | OD | 24.36 | 127,830* |
|  | JC_0905 | M | 20 | OD | 22.46 | 125,640* |
|  | JC_0914 | M | 24 | OD | 27.98 | 84,730* |
| Albinism | JC_0492 | F | 31 | OD | 23.53 | 81,810* |
|  | JC_0493 | F | 23 | OD | 22.33 | 89,120* |
|  | JC_10093 | M | 19 | OD | 21.28 | 50,400** |
|  | JC_10227 | F | 19 | OD | 22.82 | 78,890** |
|  | JC_10230 | F | 19 | OS | 20.15 | 46,020** |

\*Cone density for this subject was previously reported by Wilk et al. (2014)
\*\*Cone density for this subject was previously reported by Wilk et al. (2017)

### Fixation Testing

Fixational stability was assessed in all subjects with albinism using the fixation test module on the OPKO combined scanning laser ophthalmoscope (SLO) and optical coherence tomography (OCT) imaging system. After the SLO scanner was focused on the subject’s retina, the operator specified a group of inner retinal blood vessels to track for the duration of the run. The subject was instructed to fixate a small white cross during the run while minimizing blinks. Subjects completed three 20-second runs. Subsequently, the reference SLO images used in each run were manually registered to each other using only translation and rotation in Adobe Photoshop CS6 (Adobe Systems, Inc., San Jose, CA). The transform for each SLO image was then applied to the corresponding fixation coordinates, and the fixation points from all runs were combined to calculate the 68% and 95% bivariate contour ellipse area (BCEA) using the following equations:

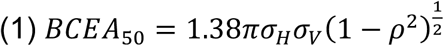

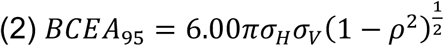

Where *σ*_H_and *σ*_V_ are the standard deviations of the coordinates in the horizontal and vertical directions (respectively) and *ρ* is product-moment correlation of the horizontal and vertical coordinates (Steinman, 1965; Crossland et al., 2004).

### Retinal Imaging

Axial length of the eye (used for lateral scaling of retinal images) was obtained for each subject using an IOL Master (Carl Zeiss Meditec, Dublin, CA). One eye for each subject was dilated and accommodation was suspended using one drop each of Phenylephrine Hydrochloride (2.5%) and Tropicamide (1%). A previously described AOSLO (Dubra and Sulai, 2011) was used to obtain images of the photoreceptor mosaic at the fovea and a strip in the temporal retina. To produce an image with minimal distortion, raw videos were first “desinusoided” to correct for the sinusoidal motion of the resonant scanner by estimating the distortion from images of a Ronchi ruling and then resampling the images over a grid of equally spaced pixels (Cooper et al., 2011). The videos were then manually inspected for reference frames that contained minimal distortion, which were then used for image registration using custom software (Dubra and Harvey, 2010). Registered images were manually aligned using Adobe Photoshop (Adobe Systems, Inc., San Jose, CA). Foveal cones were identified using a previously described, semi-automated algorithm (Garrioch et al., 2012). The location of peak cone density was identified and cone density was measured at locations across the temporal retina using a 37 μm sampling window as previously described (Wilk et al., 2014). For images within approximately 2° of the fovea, cone counting was done using the same semi-automated algorithm (Garrioch et al., 2012). For peripheral images, cones were identified manually by a single observer (MAW) using ImageJ (Schneider et al., 2012) or custom Java (Oracle Corporation, Redwood Shores, CA) software (Cooper et al., 2016).

### fMRI Visual Stimuli

All fMRI stimuli were presented on a back-projection screen mounted on the MR head coil using a BrainLogics BLMRDP-A05 MR digital projector (Puckett and DeYoe, 2015). Stimuli were generated using a ViSaGe MKII visual stimulus generator (Cambridge Research) in conjunction with MATLAB.

Stimuli included conventional expanding ring and rotating wedge retinotopic mapping stimuli (DeYoe et al., 1996). Rings and wedges were composed of black and white counterphase flickering circular checkerboards (8 Hz) with check size and ring width scaled with eccentricity. Stimuli were presented on a uniform gray background and subtended a maximum of 20° eccentricity. All subjects were instructed to continually fixate at the center of the screen. To enhance stable fixation, thin, black radial lines extending from fixation to the edge of the display were present continuously in all tasks.

To minimize unnecessary duplication, retinotopic mapping data for normal control subjects were obtained using a different experimental protocol than for the subjects with albinism. Consequently, there were slight differences in some stimulus parameters. However, due to the temporal phase mapping methods used in this study these differences did not significantly affect our results. For subjects with albinism, the wedge stimuli subtended 45° polar angle, and for control subjects the wedges subtended 90°. All subjects viewed both ring and wedge stimuli binocularly with full-field stimulation. Additionally, for subjects with albinism the expanding ring stimuli were presented to the right and left hemifields in separate runs and were tested separately for each eye. The hemifield ring stimuli were identical to the full field version, except that one hemifield was masked to match the grey background. For control subjects, the ring stimulus expanded from the center to the periphery in 40 seconds and was repeated five times per run. For subjects with albinism, both the full-field and hemifield ring stimuli expanded from 0.8° eccentricity to the periphery in 60 seconds and was repeated five times per run. For subjects with albinism, the center of the display consisted of a circular black and white disc (similar to a radioactivity symbol) with a radius of 0.8° that flickered at random intervals not synchronized to the rings/wedges presentation. To control attention, subjects were instructed to press and hold a button whenever the pattern appeared.

### fMRI Stimulus Paradigm

Control subjects completed all imaging during a single session. Subjects with albinism completed imaging during two sessions: the right eye hemifield expanding ring tasks in the first session, and all remaining tasks in the second session. All monocular hemifield runs were repeated five times and binocular full-field runs were repeated 3 times. For monocular stimuli, repetitions of the right and left hemifield stimuli were interleaved; for full-field stimuli, repetitions of the expanding ring and rotating wedge were interleaved. After each fMRI run, the subject was asked to rate their alertness on a scale from 1-5 (1 being asleep and 5 being fully awake). This measure was intended to control for subjects’ alertness, which can affect the quality of data and potentially be used as an exclusion criterion. No data were excluded from this study based on the alertness ratings.

### fMRI Acquisition

Scans were completed using a 3.0 Tesla General Electric Signa Excite 750 MRI system at the Medical College of Wisconsin. A custom 32-channel RF/Gradient head coil and a T2*-weighted gradient-echo EPI pulse sequence (TE = 25 ms, TR = 2 s, FA = 77°) were used. The 96 × 96 acquisition matrix had frequency encoding in the right-left axial plane and slice selection in the axial direction. The field of view was 240 mm and included 29 axial slices in the occipital lobe and adjacent portions of the temporal and parietal lobes with a slice thickness of 2.5 mm, yielding a raw voxel size of 2.5 mm^3^. The data were Fourier interpolated to 1.875 × 1.875 × 2.5 mm. For anatomical scans, a T1-weighted spoiled GRASS pulse sequence was used (SPGR, TE = 3.9 ms, TR = 9.6 ms, FA = 12°) with a 256 × 224 acquisition matrix. The FOV was 24 cm, and 220 slices with a slice thickness of 1.0 mm, yielding raw voxel sizes of 0.938 × 1.07 × 1.0 mm^3^. The SPGR scans were Fourier interpolated to 0.938 × 0.938 × 1.0 mm^3^ and subsequently resampled to 1.0 mm^3^. A sync pulse from the scanner at the beginning of each run triggered the onset of visual stimuli.

### Analysis Software

All fMRI data were analyzed using the AFNI/SUMA package (Cox, 1996). Surface models were produced from the high-resolution SPGR images using Freesurfer (version 5.1.0 or 5.3.0, http://surfer.nmr.mgh.harvard.edu/) with the ‘recon-all’ function.

### fMRI Pre-Processing

fMRI pre-processing was performed in the following order: reconstruction, volume registration, averaging of the time courses, removal of the initial magnetization transients, alignment. Volumes from all individual runs in an interleaved block were registered to the middle volume of the first run in the block using AFNI 3dVolreg. Individual runs for each functional task were averaged using AFNI 3dMean to produce average time courses. The before and after periods were removed using AFNI 3dcalc.

For subjects with albinism, we attempted to minimize bias in the alignment of functional scans from either of the two sessions to the reference anatomy by using a modified version of the align_across_days.csh script available on the AFNI and NIfTI server (https://sscc.nimh.nih.gov/sscc/dglen/alignmentacross2sessions). In this script, the reference SPGR anatomical images from both sessions were skull-stripped using AFNI 3dSkullStrip, aligned using AFNI align_epi_anat.py script, and averaged using AFNI 3dMean to create an average reference anatomy for the two sessions. All individual and average functional runs were then aligned to these average reference anatomies using the align_epi_anat.py script. For control subjects, all data were acquired in a single session, so creation of an average reference anatomy was unnecessary. Functional runs for control subjects were aligned to reference SPGR anatomical scans using the align_epi_anat.py script.

### Phase Encoded Retinotopic Maps

Phase encoded retinotopic activation maps were generated to plot the spatial distribution of significant fMRI responses in each of the functional tasks. Significant responses were identified by cross correlating the empirical time course data for each voxel with a reference waveform using AFNI 3ddelay (Bandettini et al., 1993; Saad et al., 2003; Datta and DeYoe, 2009). This analysis produces the correlation coefficient and phase values at the phase offset of maximum correlation for each voxel. The reference waveform used for this phase mapping procedure was a binary square wave describing the stimulus cycles convolved temporally with the “Cox Wide” estimation of the hemodynamic response function (HRF). Time courses were spatially smoothed using AFNI 3dLocalstat with a 3.75 mm spherical kernel prior to this phase mapping procedure and all functional runs were thresholded to a minimum correlation coefficient of 0.30. Phase-mapped eccentricity and polar angle values at each voxel were then projected onto functional field maps (FFmaps) using Prism View (Version 4.1.0; Prism Clinical Imaging, Elm Grove, WI) as previously described (Reitsma et al., 2013), and the eccentricity and polar angle FFmaps were manually calibrated (expanded or rotated respectively) so as to accurately represent the corresponding visual stimulus parameter. The correction factor used to make this adjustment, expressed as a constant temporal “offset” (in seconds), was then applied to the temporal delay values for all voxels in subsequent analyses.

### V1 Surface Area

Using polar angle fMRI maps on the inflated cortical surface, V1 boundaries were defined by the superior (ventrally) and inferior (dorsally) vertical meridian representations along the banks of the calcarine sulcus (**Figure 1A**). A 16° isoeccentricity boundary from the fMRI eccentricity map of V1 was marked manually on inflated cortical surfaces. This boundary was then connected to the ventral and dorsal boundaries of V1 and a region of interest (ROI) containing all nodes within this region was created. Surface area and cortical volume of V1 within this ROI was calculated using the AFNI SurfMeasures function. Surface area was calculated at both the pial surface and the gray/white matter boundary, then averaged to approximate layer 4 of striate cortex.

**Figure 1:**
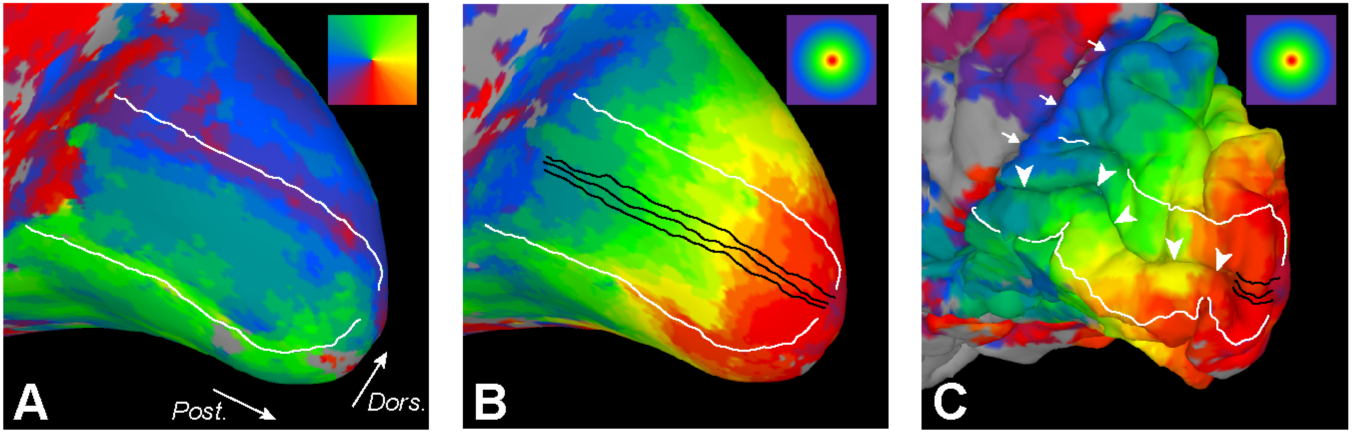
Medial occipital lobe retinotopic maps and sampling ROIs displayed on cortical surface model for a representative control subject. Color coding for polar angle and eccentricity are shown in the upper-right corner of panels A and B respectively. (A) V1/V2 boundaries (white) associated with representations of the superior and inferior vertical meridia (green and purple-red, respectively). White arrows indicate posterior (Post.) and dorsal (Dors.) orientation and apply to all panels. (B) ROIs (black) used to compute cortical mapping functions oriented parallel to the representation of the horizontal meridian within the calcarine sulcus (not shown). Phase-encoded eccentricity values (surface coloring) were assigned to each node along the linear ROIs. (C) Linear distances along the ROIs (buried within the calcarine sulcus) were calculated from the pial surface of the original folded 3D surface model (not inflated). White arrowheads indicate the calcarine sulcus. White arrows indicate parieto-occipital sulcus.

### Cortical Magnification Function Modeling

Three linear ROIs were drawn on inflated cortical surfaces along and parallel to the horizontal meridian representation within V1 for each subject (**Figure 1B**). Phase-encoded eccentricity and/or polar angle within the visual field was determined for each surface node using AFNI vol2surf and custom Matlab software. The eccentricity for each node was then matched to its linear distance along each ROI using pial (i.e., non-inflated) surface measurements in AFNI (**Figure 1C**). The total distance along each ROI was corrected by a factor of 0.93 to account for the small random variations in position of nodes in the cortical surface mesh relative to the ROIs drawn on a smooth inflated surface (correction factor determined by a previous analysis comparing mesh-based ROIs with directly-measured distances). Using custom Matlab software, cortical mapping functions were created by plotting the eccentricity represented by each voxel against its corrected distance along the ROI. Although our mapping data extended to 20°, all points beyond 16° were excluded from further analysis, because large population receptive field sizes in the periphery can introduce measurement artifacts in the shape of cortical mapping curves (Baseler et al., 2002). Additionally, in JC_10230 the ROIs from one hemisphere in one task were cropped at the beginning of the ROIs to eliminate a few highly aberrant measurements that prevented accurate curve fitting.

The cortical mapping data were then fit with a previously-described exponential curve (Engel et al., 1997), in which eccentricity in visual space (in degrees), *E*, was modeled as a function of cortical distance (in millimeters), *d*:

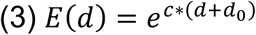

In this model, the parameter *c* is a “map scaling factor” that determines the overall shape of the function and is inversely proportional to the linear cortical magnification factor, as shown below in equation (4) (Qiu et al., 2006). Using the fminsearch function in Matlab, this curve was first fit to each individual ROI, and then normalized so that *d* was expressed as the physical distance relative to the location that represented 8° eccentricity (i.e., when *d* = 0 mm, *E*(*d*) = 8°). A single curve was then fit to all data points from all three ROIs. For all subjects, each hemisphere in each task was modeled separately.

The cortical magnification function (CMF) was subsequently computed as the derivative of the inverse of the fitted cortical mapping function, which can be represented analytically (Qiu et al., 2006) as:

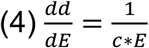

Similarly to the cortical mapping function in equation (3), here we used 1/*c* as a “CMF scaling factor” to describe the shape of the function. This curve was computed piecewise using custom Matlab software.

For comparison, we proposed a null hypothesis that CM is solely determined by cone density with no differential convergence or divergence in the connections of those cones at different eccentricities. Accordingly, the cone density for each subject was used to calculate a *predicted* cortical mapping function based on the assumption that each cone was represented by equal distance in V1. Thus, the mapping of retinal eccentricity to cortical linear distance was computed as:

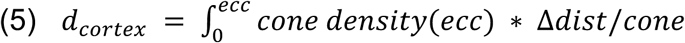

The inverse of this function provided a predicted cortical mapping function. A predicted CMF was then computed as described above for the empirical cortical mapping functions. The Δ*dist*/*cone* factor was implicitly computed so as to evenly distribute the cumulative cone count from 0 to 16 degrees over the cortical ROI distance subtending the same eccentricity range.

### Statistical Analysis

All statistical comparisons were performed using Prism 8 (GraphPad Software, Inc.). The Shapiro-Wilk test was used to assess normality, and data were classified as normal when *p* > 0.05. Parametric tests (two-tailed *t*-test and paired *t*-test) were used to compare normal data, and non-parametric tests (Mann-Whitney *U*-test and Wilcoxson matched-pairs test) were used to compare non-normal data. Differences between groups were considered to be significant when *p* < 0.05.

## RESULTS

### Fixational Stability

The 50% and 95% BCEA for each eye in each subject with albinism are shown in **Table 2**. Subjects had 50% BCEAs ranging from 0.04 to 5.97 deg^2^ with an average of 1.40 deg^2^. 95% BCEAs ranged from 0.18 to 25.97 deg^2^ with an average of 6.07 deg^2^. Previously, “steady fixation” has been defined as having 50% of fixation points fall within a 2°-diameter circle (i.e., 3.14 deg^2^ area) centered at the locus of fixation (Fujii et al., 2003). While the BCEA is not necessarily circular, its elliptical shape helps account for the predominantly horizontal eye movements that are typically observed in nystagmus. As indicated by the 50% BCEAs shown in **Table 2**, all subjects except for one (JC_10093) had 50% BCEAs that were less than 3.14 deg^2^.

**Table 2:** Fixational stability in subjects with albinism

| Subject | OD |  | OS |  |
| --- | --- | --- | --- | --- |
|  | 50% BCEA | 95% BCEA | 50% BCEA | 95% BCEA |
| JC_0492 | 0.04 | 0.19 | 0.06 | 0.28 |
| JC_0493 | 0.84 | 3.66 | 0.14 | 0.597 |
| JC_10093 | 3.68 | 16.01 | 5.97 | 25.95 |
| JC_10227 | 0.24 | 1.05 | 0.45 | 1.95 |
| JC_10230 | 1.92 | 8.35 | 0.62 | 2.68 |

OD = right eye; OS = left eye; BCEA = bivariate contour ellipse area. All values are expressed in degrees<sup>2</sup>.

### Cone Density

Control subjects’ foveal peak cone densities ranged from 84,733 to 165,080 cones/mm^2^ (mean ± SD: 123,885 ± 28,760 cones/mm^2^), while patients with albinism had foveal cone densities ranging from 46,019 to 89,116 cones/mm^2^ (average ± SD = 69,248 ± 19,623 cones/mm^2^). Cone density as a function of eccentricity for each subject can be seen in **Figure 2**. All subjects showed a reduction in cone density with eccentricity. Patients with albinism had generally lower densities within 3 degrees of the fovea; however, cone density became more similar across all subjects in the periphery (**Figure 2**). Three subjects with albinism (JC_0492, JC_0493, and JC_10227) had peak cone densities that fell within two standard deviations of normal, while the remaining subjects with albinism (JC_10093 and JC_10230) had much lower peak densities.

**Figure 2:**
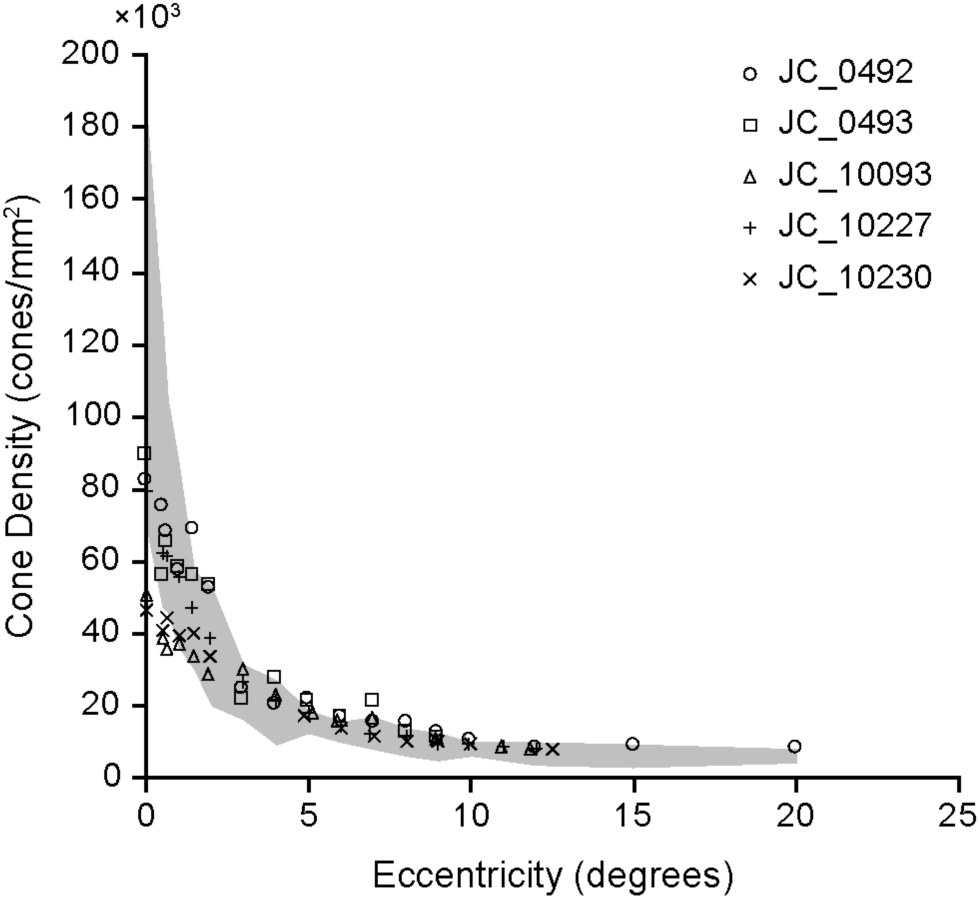
Retinal cone density as a function of eccentricity. Each subject with albinism is represented by a different data point symbol. Gray shaded area represents the average for all control subjects ± 2 SD.

### Surface Area of V1

The cortical surface area and volume of V1 representing the central 16° of the visual field was measured in all control subjects and four subjects with albinism using manually-drawn ROIs based on isopolar and isoeccentricity maps (**Figure 3A**). V1 was not measured in JC_10230 because the thresholded eccentricity data (i.e., voxels with a correlation coefficient of 0.30 or greater) did not extend all the way to 16°. While surface area appeared to be reduced in albinism relative to controls (**Figure 3B**), this difference approached but did not achieve statistical significance (two-tailed *t* test: *t* = 1.98, df = 16, *p* = 0.065).

**Figure 3:**
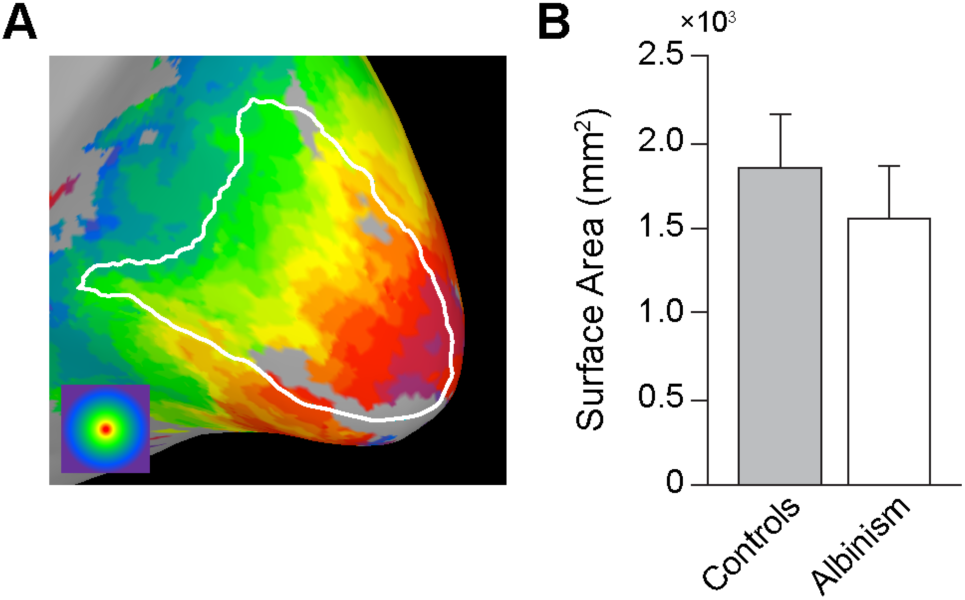
Mean surface area and gray matter volume of V1 is not significantly different for albinism and control subjects (two-tailed t test: t = 1.98, df = 16, p = 0.065). Boundaries used to compute V1 shown in Figure 1. Error bars represent one standard deviation. Values for subjects with albinism are from only four subjects due to incompleteness of cortical maps in one subject (JC_10230).

### Retinotopy in Albinism

For control subjects, isoeccentricity retinotopic maps were obtained using binocular, full-field stimuli (**Figure 4A**), which showed normal retinotopic organization in V1 (**Figure 4B**). For subjects with albinism, retinotopic maps (**Figure 4C**) were acquired using monocular hemifield stimuli (**Figure 4A**) as described previously (Hoffmann et al., 2003). In all subjects, the most complete and contiguous retinotopic activation was produced in the hemisphere contralateral to the visual stimulus, as would normally be expected (**Figure 4C**, columns 1 and 4; see also data for the left eye in Supplemental Figure S1). However, each hemisphere could also be activated by the ipsilateral visual field, which resulted in overlaid representations of both hemifields within the same hemisphere, a highly aberrant result (**Figure 4C**, columns 1 and 3 for the left hemisphere, columns 2 and 4 for right hemisphere). These aberrant hemifield representations were asymmetric, with the most extensive activation in the hemisphere contralateral to the stimulated eye. Thus, when the right eye was stimulated (as in **Figure 4C**) the aberrant activation of the left hemisphere was more prominent than that in the right hemisphere (**Figure 4C**, column 3 greater than column 2), and when the left eye was stimulated the aberrant activation of the right hemisphere was more prominent than that in the left hemisphere (Supplemental Figure S1, column 2 greater than column 3). This asymmetry is also evident when comparing the normal activation to the aberrant activation in the same hemisphere: for example, in subject JC_0492, when the right eye was stimulated the normal and aberrant representations in the left hemisphere (i.e., contralateral hemisphere; **Figure 4C**, columns 1 and 3) appeared more similar than the two representations in the right hemisphere (i.e., ipsilateral hemisphere; **Figure 4C**, columns 2 and 4). This was a general trend across subjects with albinism. There was a similar trend for the left eye stimulus, with the greatest similarity between normal and aberrant representations in the right hemisphere (see Supplemental Figure S1, where columns 2 and 4 were more similar to each other than columns 1 and 3).

**Figure 4:**
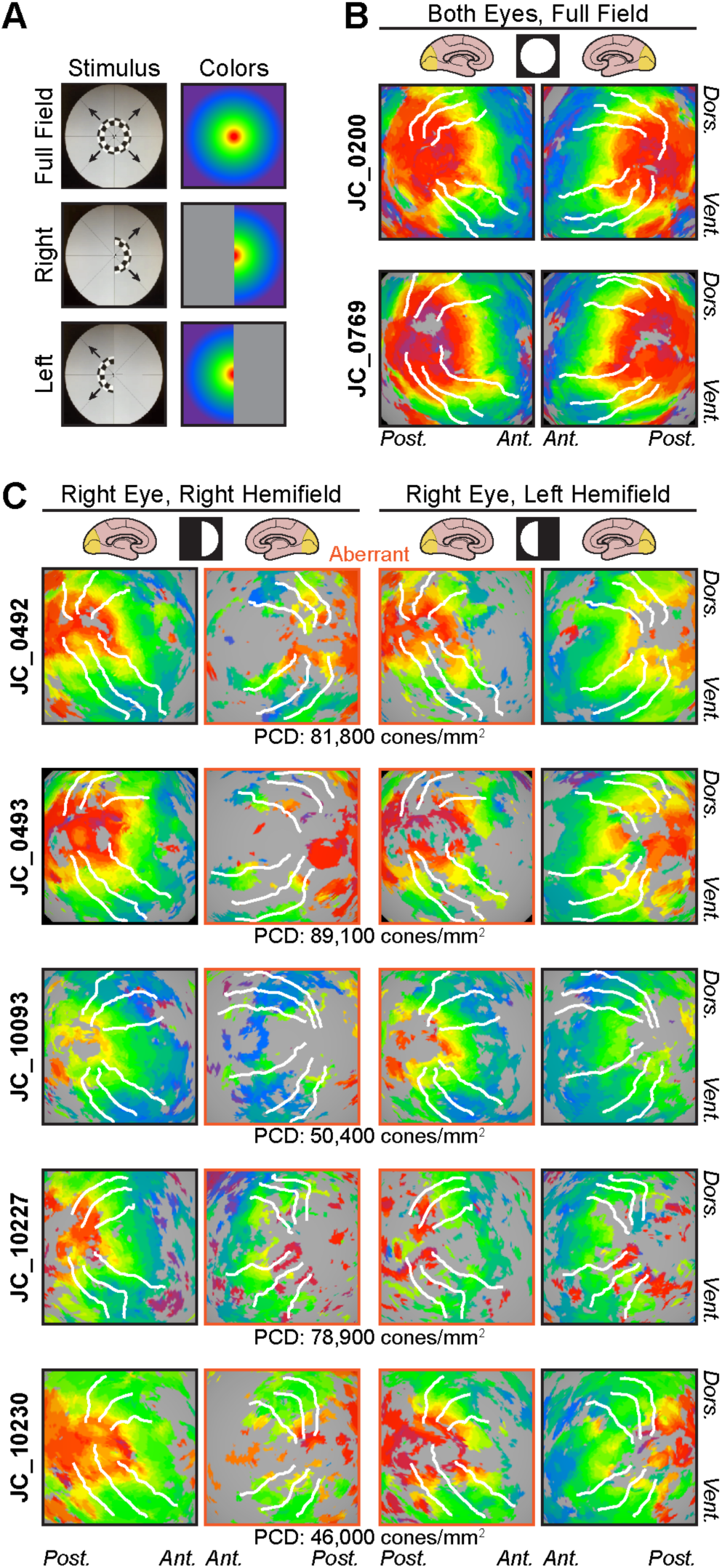
Retinotopic maps of visual field eccentricity. (A) Visual stimuli and color coding used for eccentricity mapping in controls (full field) and subjects with albinism (hemifield stimuli). (B) Eccentricity maps for two representative control subjects. All control subjects were tested using binocular viewing of the full-field ring stimulus. (C) Eccentricity maps for subjects with albinism, obtained using hemifield ring stimuli viewed monocularly. Data for right eye viewing condition are shown here (see Supplemental Figure S1 for left eye data). Retinotopy patterns outlined in orange (middle columns) are aberrant ipsilateral hemifield representations. Peak cone densities for each subject with albinism are indicated below each row. All retinotopic maps are displayed on spherically-inflated cortical surface models. Visual stimuli are indicated by white circle/semicircle symbols at head of respective columns. White lines in B, C mark dorsal and ventral boundaries of V1/2/3 based on polar angle data (cf. Figure 1). PCD = peak cone density; Dors. = dorsal; Vent. = ventral; Ant. = anterior; Post. = posterior.

In addition to these trends, however, there was also significant variation across our albinism cohort (compare down each column of **Figure 4C**), which was particularly evident with respect to the overall continuity of the isoeccentricity maps. This can be seen clearly in subjects JC_0492 and JC_10227, who showed some of the most and least contiguous maps: JC_0492 had a nearly complete retinotopic representation with few holes (especially in the left hemisphere), while JC_10227 showed only sparse activation in either hemisphere when the right visual field was stimulated. Additionally, JC_10227 showed notable activation by the peripheral visual field in the ipsilateral hemisphere regardless of which hemifield was stimulated, which was not as prominent in other subjects (**Figure 4C**, second column).

Subjects in our albinism cohort also varied considerably in the predominance of central versus peripheral field representations (note dominant colors in each subject in **Figure 4C**). This is illustrated by directly comparing subjects JC_0493 and JC_10093: JC_0493 had much more central (red/orange) activation than JC_10093, but JC_10093 had greater peripheral (blue) activation than JC_0493. Additionally, subjects varied in the spatial localization of the isoeccentricity bands: subjects who had weak foveal activation also had peripheral activation that was apparent nearer to the occipital pole, where the foveal confluence is typically found (compare the location of the yellow band across subjects in **Figure 4C**). In general, the extent of activation by the central visual field appeared to correlate with subjects’ peak cone densities (shown below each row of images in **Figure 4C**), but JC_10230 was a notable exception. This subject had the lowest peak cone density but showed much clearer activation by central visual field than JC_10093, who had the next-lowest peak cone density.

### Cortical Magnification in Albinism

Cortical mapping in V1 was then measured quantitatively. Examples of cortical mapping functions for representative controls and subjects with albinism are shown in **Figure 5A-B** (control) and **Figure 5E-F** (albinism). The empirical data are shown by the red and blue points for the right and left hemispheres, respectively (and for the 3 ROIs within each hemisphere). Fitted curves (equation 3, Methods) are shown as the smooth colored lines (See Supplemental Figure S2 for data/curves for all subjects). The fitted curves for all subjects are shown in **Figure 5C** and **5G**. When the map scaling factor, *c*, from the fitted cortical mapping functions was compared across hemispheres, there was no significant difference between the right and left hemispheres (Wilcoxson matched-pairs test, *p* > 0.99).

**Figure 5:**
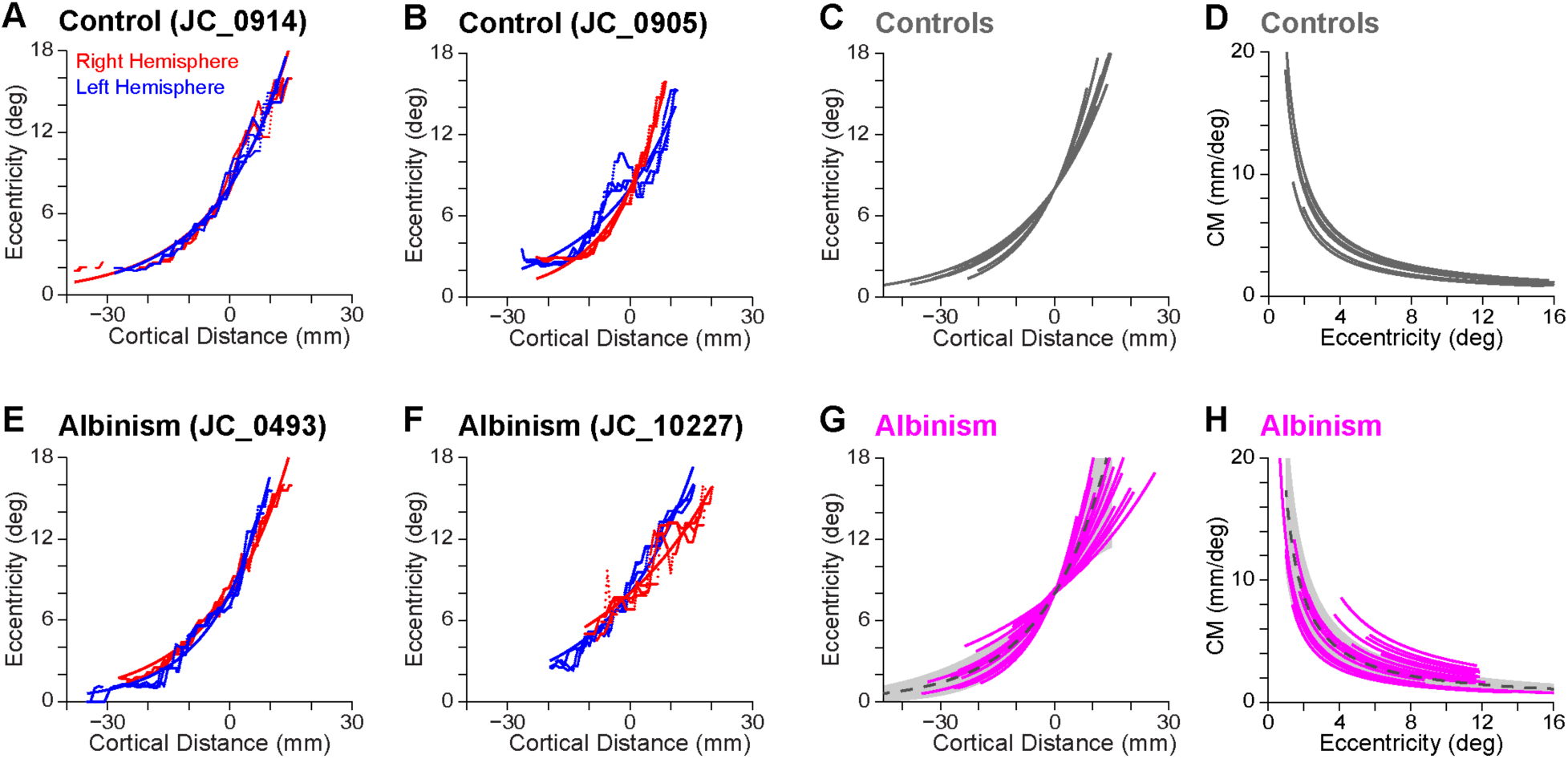
Cortical mapping and magnification functions in controls and subjects with albinism. (A-B) Cortical mapping data and fitted functions for two representative control subjects, including data from the left (blue) and right (red) hemispheres. (E-F) Cortical mapping data and fitted functions for two representative subjects with albinism. Data for left hemisphere (blue) obtained with right eye, right hemifield stimulation. Data for right hemisphere (red) obtained with left eye, left hemifield stimulation. (C) Fitted cortical mapping functions and (D) corresponding CMFs (see Methods) for both hemispheres of all control subjects (gray curves). (G) Fitted cortical mapping functions and (H) corresponding CMFs for subjects with albinism (magenta curves). Dashed dark gray lines and gray shaded regions show mean ± 1 SD for controls. In G and H data are combined for left hemisphere, right hemifield stimulus and right hemisphere, left hemifield stimulus and for both right and left eye viewing conditions.

In subjects with albinism, each visual hemifield in each eye was stimulated separately, resulting in eight possible cortical mapping functions for each subject (2 hemispheres x 2 visual hemifields x 2 eyes). However, due to the limited extent of activation by the ipsilateral hemifield in either hemisphere, **Figure 5E-H** shows only the curves representing activation by the contralateral visual hemifield. Since each eye was stimulated separately, the cortical mapping functions from each eye were compared within each hemisphere by comparing the map scaling factors (*c* from equation (3)). The functions from the right and left eyes were not significantly different (paired *t*-test: *t* = 0.24, df = 9, *p* = 0.82), and the functions from the same eye were not significantly different between hemispheres (paired *t*-test: *t* = 0.42, df = 9, *p* = 0.68). Mapping functions were also compared between controls and subjects with albinism (**Figure 5C** versus **5G**). When the map scaling factors for all control subject hemispheres (mean ± SD: 0.0587 ± 0.0086) were compared to those for subjects with albinism (mean ± SD: 0.0566 ± 0.0188), there was no significant difference (Mann-Whitney *U*-test: *U* = 82, *p* = 0.44).

The size of V1 is known to directly impact the magnitude of estimated CM, though the shape of the function does not change significantly (Sereno et al., 1995). In order to verify that the trend toward smaller V1 surface area in subjects with albinism did not also affect the above comparison between control and albinism groups, the map scaling factor in each task was compared to the V1 surface area of the corresponding hemisphere. The map scaling factors were not significantly correlated with V1 surface area in either control (*r*^2^= 0.22, *p* = 0.17) or albinism groups (right eye: *r*^2^= 0.27, *p* = 0.19; left eye: *r*^2^= 0.28, *p* = 0.17).

The fitted cortical mapping functions were then used to derive the cortical magnification function (CMF) for each task in each subject (see Equation (4) in Methods). The resulting functions for control and albinism subjects are shown in **Figure 5D** and **5H** respectively. CMFs for control subjects were modeled separately for each hemisphere. For subjects with albinism, CMFs were modeled separately for each eye and each hemisphere (stimulated by the contralateral visual field). When the CMF scaling factors were compared between control subjects (mean ± SD: 17.33 ± 2.26) and subjects with albinism (mean ± SD: 19.75 ± 6.93), there was no significant difference (Mann-Whitney *U*-test: *U* = 82, *p* = 0.44). However, the CMF scaling factor was more variable in albinism relative to controls (*F* test: *F* = 9.45, DFn = 19, Dfd = 9, *p* = 0.002). This is evident in **Figure 5D** versus **5H** (in 5H, the dashed gray line is the average of the fitted curves from control subjects, and the shaded gray area represents the values within 1 SD of the average).

As mentioned above, all subjects with albinism had aberrant ipsilateral hemifield representations superimposed on the normal representation of the contralateral visual field (**Figure 4C**). In most subjects this ipsilateral activation was highly truncated compared to the contralateral activation. However, two subjects had at least one hemisphere with sufficient ipsilateral activation to compare the aberrant activation to the “normal” activation within the same hemisphere. **Figure 6** shows the comparison between the aberrant ipsilateral field mapping data (red) and the normal contralateral field mapping data (gray) in these unique subjects. In JC_10093, the data from the two hemifields did not match, but rather the aberrant activation showed a unique pattern compared to the normal activation. When only the contralateral hemifield was stimulated, activation in the same hemisphere but from different eyes showed much better agreement (see Supplemental Figure S2 for comparison). In JC_10227, the aberrant activation appeared to agree with the normal activation overall. However, in the aberrant activation there was notable variability between ROIs, particularly in the peripheral visual field, which was not as pronounced in the normal activation.

**Figure 6:**
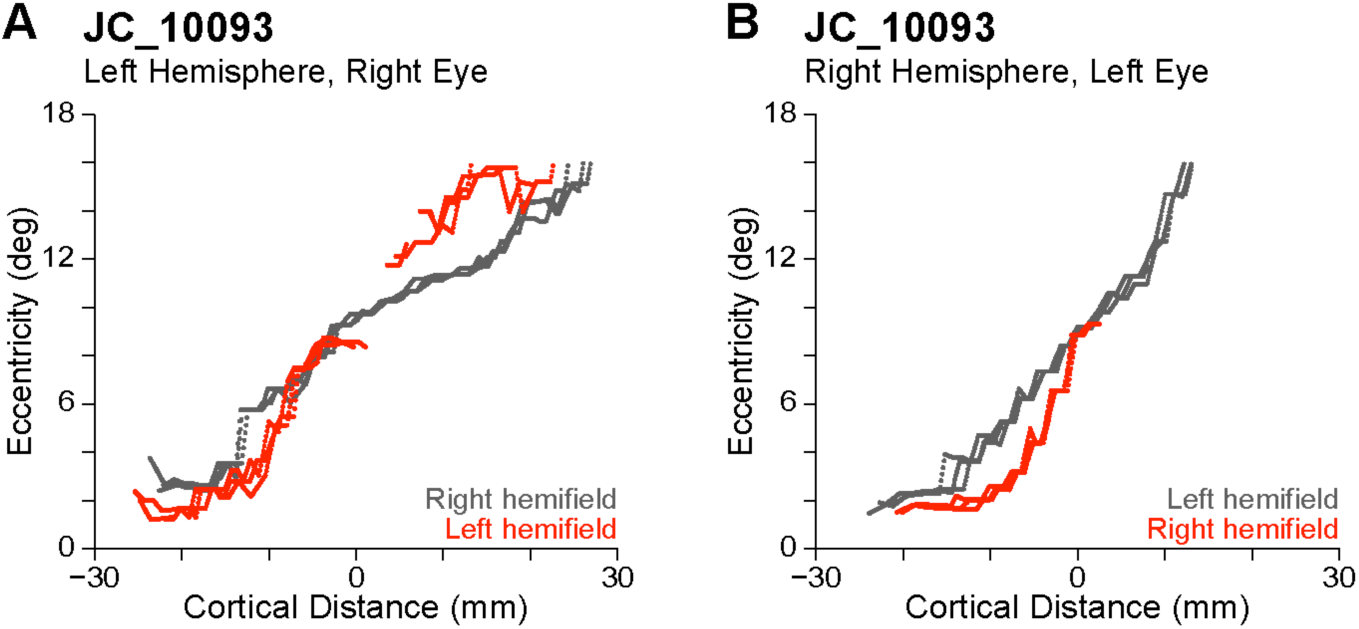
Comparison of empirical cortical mapping functions for normal contralateral (gray) versus aberrant ipsilateral (red) hemifield stimulation in left and right hemispheres of subject JC_10093. The red and gray data are from the same physical ROIs yet are distinctly different. LH = left hemisphere; RH = right hemisphere.

### Empirically-Based vs. Cone-Predicted Models of Cortical Magnification

**Figure 7** shows empirically-derived CMFs (red) for each subject with albinism along with a predicted CMF (green) based on cone density alone. Control results are shown in **Figure 8.** As outlined in Methods, the cone density predictions have been intentionally aligned to the empirically-based curves at 8° eccentricity, to facilitate comparison of the *shapes* of the curves rather than their absolute differences (which are sensitive to an arbitrary scaling factor). It is clear in all hemispheres for which the data extend to within 2° of the center of gaze (**Figure 7**, dark gray shaded region) that the empirically measured CM increases far more rapidly than is predicted by cone density alone, and this is also the case (albeit to a lesser extent) between 2-4° eccentricity (**Figure 7**, light gray shaded region). This is true for both albinism (**Figure 7**) and control groups (**Figure 8**).

**Figure 7:**
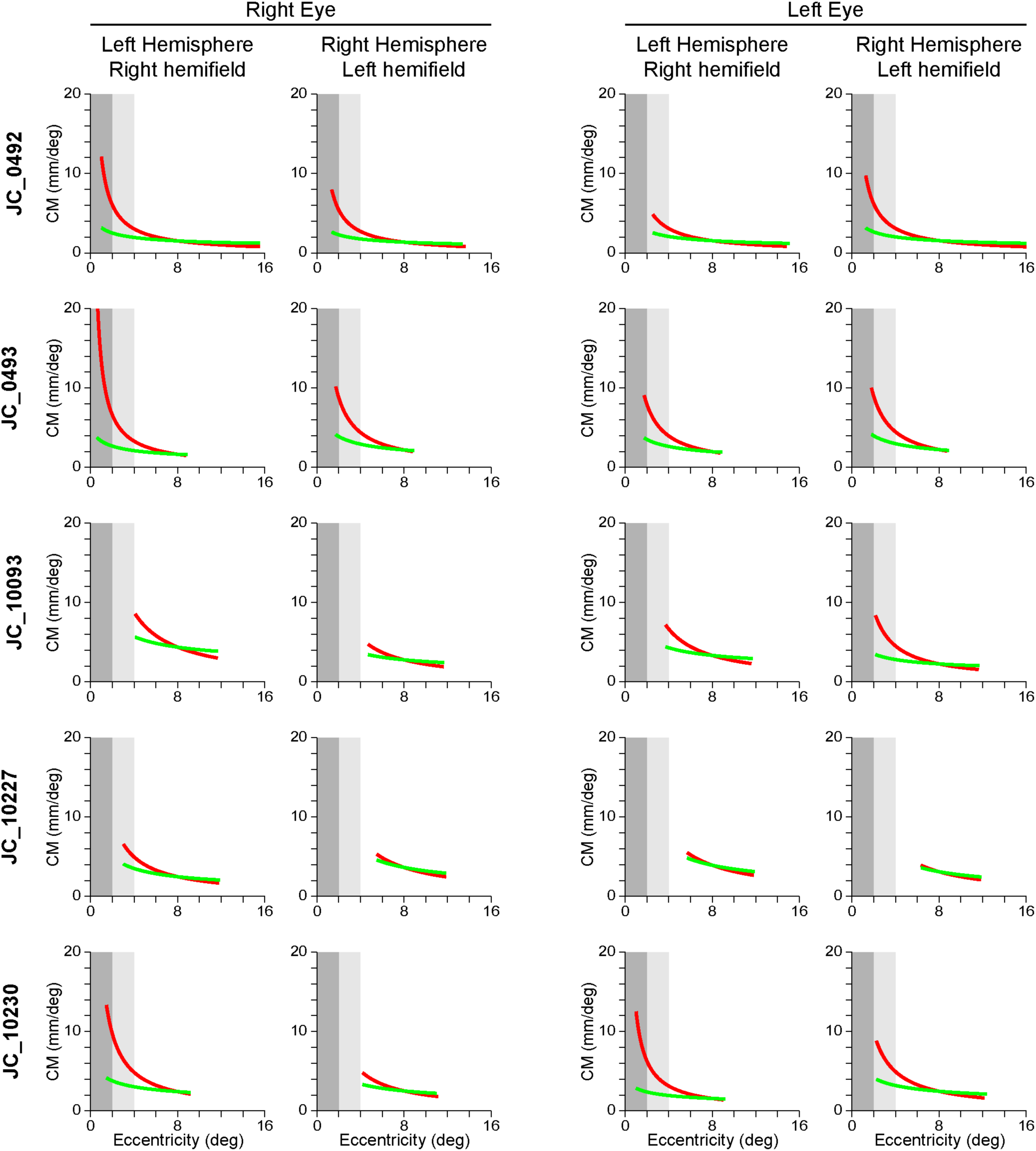
Cortical magnification functions based on empirical data (red) and cone density predictions (green) for subjects with albinism. To aid comparison, eccentricity ranges from 0-2° and 2-4° are shaded in dark and light gray respectively.

**Figure 8:**
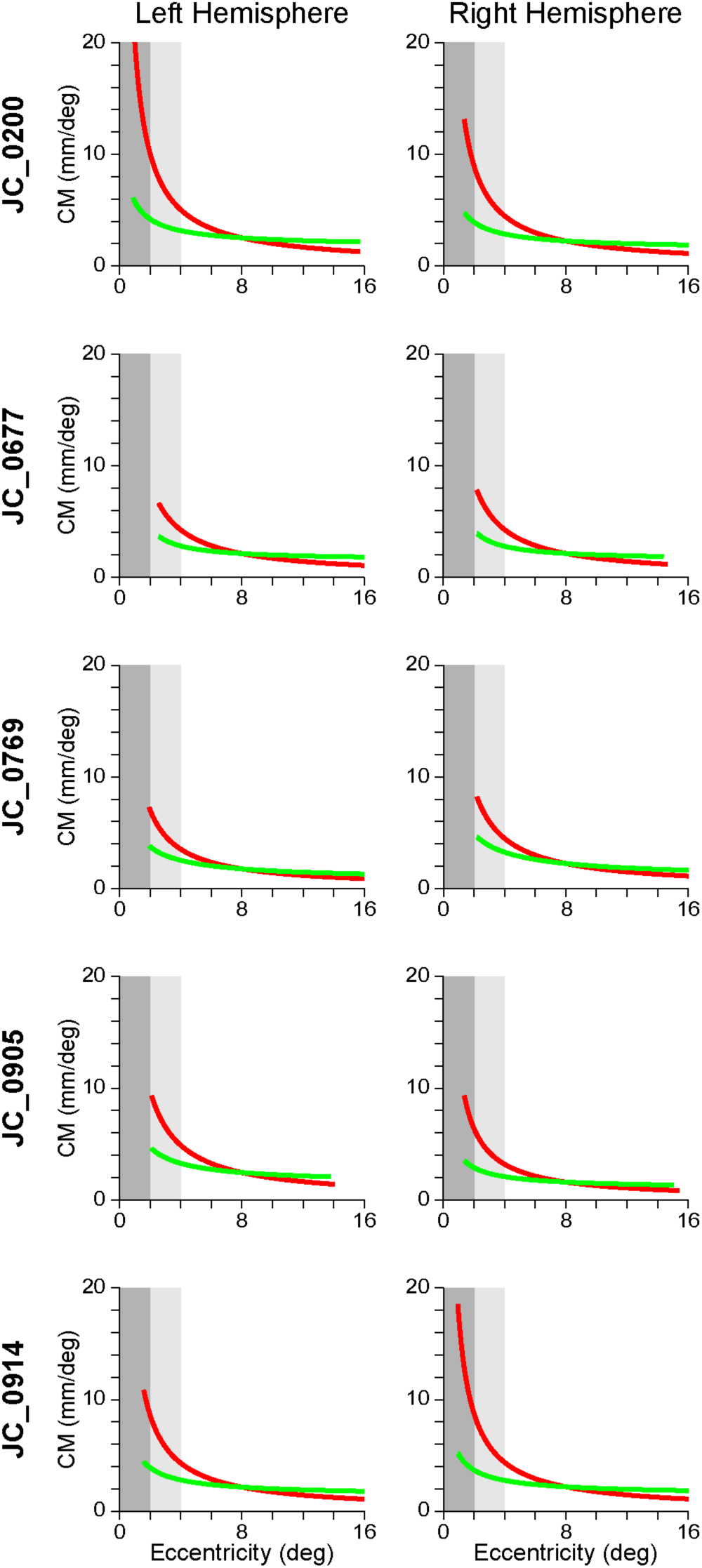
Cortical magnification functions based on empirical data (red) and cone density predictions (green) for control subjects. To aid comparison, eccentricity ranges from 0-2° and 2-4° are shaded in dark and light gray respectively.

In order to determine whether cortical organization was correlated with cone density, the cortical mapping functions and CMFs were also compared to each subject’s peak cone density. Since cone density was measured in only one eye in each subject, only the cortical functions based on activation by the imaged eye (i.e., the right eye in all subjects except JC_10230; see **Table 1**) was included in this analysis. The average map scaling factor (*c*) and the CMF scaling factor (1/*c*) from both hemispheres was plotted against peak cone density for each subject. Neither the map scaling factor nor the CMF scaling factor was significantly correlated with peak cone density in either control subjects (*c*: *r*^2^= 0.053, *p* = 0.71; 1/*c*: *r*^2^= 0.067, *p* = 0.67) or in subjects with albinism (*c*: *r*^2^= 0.35, *p* = 0.29; 1/*c*: *r*^2^= 0.17, *p* = 0.49).

## DISCUSSION

### Aberrant Retinotopy in Albinism

The results of this study are largely consistent with previous reports of highly aberrant retinotopic organization in visual area V1 of subjects with albinism (Creel et al., 1974; Dorey et al., 2003; Hoffmann et al., 2003; Schmitz et al., 2004) but significantly extend those findings to include an analysis of individual variability and a quantitative account of CM compared with individual retinal cone density measurements. Aberrant retinotopic organization was apparent in all subjects with albinism and consisted of superimposed representations of opposite hemifield representations within the same hemisphere. The aberrant (ipsilateral) hemifield activation was most prominent in the hemisphere contralateral to the eye being stimulated. This aberrant activation typically represented regions of the visual field within approximately 5-6° of the center of gaze but varied considerably across the albinism group. This agrees with previous studies that found significant variation in the left-right margin of aberrant decussation among individuals with albinism (Dorey et al., 2003; Hoffmann et al., 2005; von dem Hagen et al., 2007). Overall, these findings support previous descriptions of pathological optic nerve decussation at the optic chiasm in albinism. Ganglion cell projections from the temporal retina that ordinarily synapse in the ipsilateral hemisphere instead decussate to the contralateral hemisphere (Creel et al., 1978; Hoffmann et al., 2003), giving rise to the aberrant cortical retinotopy.

When the surface area of V1 was measured in this study, subjects with albinism had modestly reduced surface area relative to control subjects, but this difference was not statistically significant (**Figure 3**). This is not surprising, because changes in occipital gray matter volume that were reported in previous studies of albinism were highly localized to the occipital pole (von dem Hagen et al., 2005; Bridge et al., 2014); therefore, measurements of V1 surface area as a whole are less likely to be dramatically affected. However, subjects with albinism are also known to have reduced optic nerve, optic chiasm, and optic tract size (von dem Hagen et al., 2005; Ather et al., 2019), which may reflect a reduction in retinal afferents from the foveal region where cone density is reduced (Wilk et al., 2014). In normal visual system development V1 surface area is thought to be correlated with optic tract size (Andrews et al., 1997). Thus, the apparent lack of correlation between previous reports of reduced optic tract size in albinism and our measurements of nearly normal V1 surface area in albinism suggests that unique factors involved in the development of retinocortical projections in albinism may preserve cortical space despite a reduced number of retinal afferents.

### Variability in Retinotopic Organization in Albinism

In the albinism group, we noted marked variations in features of the overall pattern of retinotopic organization that have not been appreciated previously. These features included both the continuity of the retinotopic maps and the relative surface area devoted to central versus peripheral visual field activation. Such variability within the albinism population is consistent with previous studies that have shown significant variability in retinal phenotypes among individuals with albinism, particularly in the severity of foveal hypoplasia (Thomas et al., 2011; Kruijt et al., 2018) and in peak cone packing density (Wilk et al., 2014; Wilk et al., 2017). Indeed, our group previously performed a detailed analysis of the retinal structure of the subjects presented in this study (Wilk et al., 2014; Wilk et al., 2017). The results of that study were used to intentionally select subjects with albinism for fMRI imaging who represented a broad spectrum of retinal structure (for cone densities, see **Figure 2**).

Overall, the area of cortical activation corresponding to the central visual field in subjects with albinism appeared to be reduced compared to control subjects. This may be partly due to the visual task used for retinotopic mapping in subjects with albinism, because the smallest annulus in the expanding ring stimulus did not include the foveal center. Rather, the center of fixation was covered by the round fixation marker that had a radius of 0.8° and appeared randomly to control for attention. While the central activation might have been improved by using a contracting ring or drifting bar stimulus (Dumoulin and Wandell, 2008), the presence of robust foveal activation in some subjects with albinism (e.g. JC_0492 and JC_0493) indicates that the stimulus itself was not sufficient to prevent strong activation by central regions of the visual field. It is also possible that variation in the continuity of the retinotopic maps could be due to reduced subject alertness during imaging, so we explicitly monitored alertness by self-report after every fMRI scan. Only one subject (JC_0493) reported alertness levels below 3 (on a scale from 1 to 5, where 5 is most alert), yet this subject had some of the most robust activation patterns. Therefore, we do not believe the reduced foveal representation in albinism to be artifactual. Moreover, this observation is consistent with previous fMRI-based findings in albinism (Schmitz et al., 2004). As mentioned above, known structural changes in V1 in albinism are also preferentially found at the occipital pole (von dem Hagen et al., 2005; Bridge et al., 2014), which further corroborates an overall reduction in the functional activation of the foveal region.

We also found that the decreased foveal activation correlated with peak cone density in four of five subjects with albinism. However, the presence of one subject who did not fit this trend (JC_10230) indicates that factors other than cone density (for example, the divergence or convergence of retino-cortical projections) may also contribute to the relative sizes of the central and peripheral retinotopic representations.

The extent of central field activation across subjects with albinism tended to be inversely correlated with the extent of peripheral field activation. That is, in subjects who had reduced foveal activation the peripheral regions of the visual field tended to expand toward the occipital pole (see **Figures 4C and S1**, columns 1 and 4). One possible explanation for this phenomenon is that, when input from the central visual field is reduced or absent during development, retino-cortical afferents representing more peripheral regions of the visual field may expand into cortical space normally allocated to the more central portions of the visual field. This would also explain our finding that V1 surface area is relatively normal despite a known reduction in retinal afferents that is concentrated at the fovea (Wilk et al., 2014) and in optic tract size (von dem Hagen et al., 2005; Ather et al., 2019). A similar cortical expansion of non-foveal afferents has been described previously in achromatopsia. In this genetic disease there is a complete absence of functional cone photoreceptors, yet stimulation of rod photoreceptors leads to aberrant cortical activation near the occipital pole (Baseler et al., 2002). Thus, it is possible that analogous cortical reorganization may occur in albinism, in which peripheral inputs “spread out” into cortical regions that might otherwise be occupied by foveal afferents. A second possible explanation is that a release of lateral inhibition by the visual cortical neurons that are typically activated by the central visual field allows the neighboring retinotopic areas to become more active, though it is unclear how such an effect might propagate throughout the map to permit expansion of the most peripheral representation.

### Cortical Magnification: Albinism vs. Controls

Despite grossly aberrant hemifield organization, the cortical mapping functions for subjects with albinism were *not* (on average) significantly different from those for control subjects (**Figure 5G**; albinism curves shown in orange, average control function shown in gray). Our control data also match those of Sereno et al. and Engel et al. (Sereno et al., 1995; Engel et al., 1997). This similarity between control subjects and subjects with albinism was also evident when the fitted cortical mapping functions were used to model the CMF in each hemisphere (**Figure 5H**). This allows us to reject our initial hypothesis that reduced cone density in albinism would necessarily lead to reduced CM. Instead, some of the hemispheres of subjects with albinism tended to have *greater* CM in peripheral regions of the visual field compared to the control group (**Figure 5H**, orange curves outside the gray underlay representing controls). This trend in CM would not be explained by overall surface area of V1, because (as discussed above) the surface area of V1 was not significantly different in albinism than in controls, and in fact trended toward *decreased* V1 surface area. This suggests that, in albinism, retinal cone density does not determine the size of the retinotopic representation in V1 as has previously been suggested (Dougherty et al., 2003), but rather that the cones that are present (along with their downstream synaptic partners) capitalize on the entirety of the cortical space available to them. In this scenario, it is possible that the amount of cortical space devoted to V1 (and other visual areas) is primarily determined by non-retinal factors, which may be unique to albinism (or similar genetic conditions).

It is also notable that, for one of the subjects with albinism (e.g. JC_10093), the cortical eccentricity mapping functions for the normal and aberrant fields had local zones in which the eccentricities differed markedly despite being represented by exactly the same cortical voxels (**Figure 6**). This indicates that the two hemifield representations are not always precisely in mirror-image register, and that more subtle wiring anomalies can occur in addition to those related to a shift in the line of left-right decussation at the optic chiasm. Though one might suppose that these more subtle errors simply represent random variations, it is important to appreciate the precision of same hemifield overlap in normal controls that gives rise to the systematic and precise computations of retinal disparity responsible for stereopsis. Whether the lack of precise registration between opposing hemifield representations in albinism is functionally important remains unclear.

### Cortical Magnification: Empirical vs. Cone-Based Predictions

The relationship between CM and cone density was explored further by comparing the empirically-derived CMF models with a predicted CMF model based on cone density alone, assuming that each cone was allocated an equal amount of cortical space in V1. All subjects—both controls and albinism—showed a greater increase in CM near the center of the visual field than would be predicted based on cone density alone (**Figures 7**, **8**). When the empirical and cone-predicted CMFs are compared in the central-most regions (see shaded areas in **Figure 7**), it is evident that both the magnitude and slope of the empirical functions (red) are greater than the cone-based predictions (green). Note, however, that the empirical data within the central 2 degrees (dark gray zone) are marginal due to limitations of the stimulus (see Methods), and this was particularly true for two of our subjects with albinism (JC_10093, JC_10227). This is important because one might expect that the extreme loss in cone density observed in albinism relative to control subjects would lead to markedly reduced CMFs. However, the biggest differences in cone density are limited to the fovea and don’t extend beyond 2° where our more reliable CM data begin. This might also account for the failure to observe major differences in CMF between subjects with albinism and controls.

That empirical CM values exceed the cone-based predictions might be explained by a number of factors. First, it is known that there is greater convergence of cones onto peripheral retinal ganglion cells (RGCs) compared to central RGCs (Curcio and Allen, 1990; Dacey and Petersen, 1992; Dacey, 1993), which would exacerbate cone-based differences in central versus peripheral magnification. Moreover, RGCs with receptive fields near the center of the visual field are thought to project to more cortical space than peripheral RGCs (Azzopardi and Cowey, 1996; Popovic and Sjöstrand, 2001), though this is debated (Wässle et al., 1990). In order to examine the representation of RGCs in V1, one must consider both the convergence/divergence of GC projections onto LGN neurons and divergence of LGN projections on to layer 4 of V1. Indeed, Azzopardi and Cowey (Azzopardi and Cowey, 1996) argue that the final V1 cortical magnification function is the result of successive expansion of the foveal representation both in the LGN and in the cortex. The stage between the LGN and cortex was described in detail by Connolly and Van Essen (Connolly and Van Essen, 1984) for the macaque monkey. They found that “*The total number of cortical neurons per LGN neuron is about 130 on average, but it extends over approximately a tenfold range, from less than 100 in the far periphery to nearly 1,000 in the fovea.*” Whether this tenfold difference in divergence is also true for humans with or without albinism is unknown, but is likely to be a major factor in determining the resulting cortical magnification. Clearly, a comprehensive, quantitative account of the neural basis of CMF must await more extensive estimates of convergence/divergence versus eccentricity at each stage of the retinostriate hierarchy.

### Behavioral Predictions

Given previous observations that human visual acuity is normally correlated with CM (Duncan and Boynton, 2003), the similarities in CMFs between controls and subjects with albinism in this study might predict that parafoveal and peripheral visual acuity would be similar in albinism to that in normal controls. A previous study of acuity in albinism found that central visual acuity in albinism was reduced but peripheral visual acuity was similar to normal controls, which supports this prediction (Wilson et al., 1988). However, that study only measured acuity at the center of gaze and at 10° inferior, so it is currently unknown at what point between the fovea and 10° the acuity thresholds in albinism approach normal levels. While our measurements of CM were limited to 2-16° (i.e., they did not extend to the fovea), the empirically-derived CMFs in subjects who had the most extensive central representations appear to remain within normal limits (see **Figure 5H**). If this trend continues all the way to the fovea, it would indicate that the CMFs may not necessarily account for the reduced central visual acuity typically observed in albinism. However, the variability in cortical organization shown in this study indicates a role for plasticity in modifying the functional relationship between retinal and cortical structures. More detailed studies of visual acuity are needed in this population along with targeted investigations of the cortical foveal confluence in order to further explore the etiology of the visual acuity deficits in albinism (Azzopardi and Cowey, 1996).

### Limitations

One of the primary limitations of this study is the small sample size. Both the rarity of the disease and the prevalence of moderate to severe nystagmus within this population serve as barriers to recruitment of large numbers of subjects with albinism who are good candidates for fMRI. However, the finding that there is significant variability in retinotopic organization between individuals with albinism only increases the importance of measuring cortical phenotypes in more individuals to determine the full extent of phenotypic variability. While we aim to increase our sample size in the future, the current small cohort precludes observation of strong correlations between genetic subtypes of albinism and distinct cortical phenotypes. Learning more about such genotype/phenotype correlations in albinism will be essential for guiding clinical diagnosis and for developing interventional therapies.

An important methodological limitation of the current study is the use of expanding checkered annuli to assess the eccentricity dimension of cortical retinotopy. The presence of a relatively large fixation marker and the limited resolution of our visual stimulus display within the MRI scanner compromised our ability to obtain reliable mapping data with approximately 1.5 degrees of fixation. Future use of drifting bar stimuli (Dumoulin and Wandell, 2008) in conjunction with an improved video display and fixation marker should ameliorate this problem.

Another potential concern is that nystagmus and eccentric fixation are common in albinism and might adversely affect the retinotopic maps. While nystagmus can introduce noise into the fMRI signal, it is less likely to systematically alter the spatial properties of a retinotopic map (Baseler et al., 2002). Moreover, BCEA values comparable to those reported here for all but one subject with albinism do not appear to be correlated with population receptive field (pRF) sizes in other populations with unsteady fixation (Clavagnier et al., 2015). Eccentric fixation, on the other hand, can affect the shape and symmetry of cortical mapping functions (Baseler et al., 2002). However, the cortical mapping functions that we observed tended to be notably symmetric between hemispheres suggesting that eccentric fixation is unlikely to have had a significant effect on the data presented here.

## CONCLUSIONS

This study both confirms previous findings of abnormal retinotopic organization in albinism and expands upon these findings by showing that there is greater diversity in retinotopy between individuals with albinism than has previously been appreciated. While these changes often correlate with retinal cone density, this is not always the case. Cortical magnification outside the fovea is not significantly different in albinism than in normal controls and is greater in both groups than is predicted by cone density alone. This indicates that post-receptoral mechanisms responsible for CM in albinism are at least qualitatively, if not quantitatively, similar to those in normal controls.

Overall, our results show that the pattern of retinocortical miswiring that has previously been ascribed to aberrant left-right decussation at the optic chiasm is significantly more complex and varied than previously thought. Whether this additional complexity occurs at the optic chiasm or represents additional connectivity changes downstream is unclear. Albinism provides an excellent model in which both peripheral and central effects of genetic mutation can be explored quantitatively.

## ACKNOWLEDGEMENTS

The authors would like to thank Brittany Bartlein, Robert Cooper, and Phyllis Summerfelt for their contributions to this work. Research reported in this publication was supported by the National Eye Institute, the National Institute of General Medical Sciences, and the National Center for Advancing Translational Sciences of the National Institutes of Health under award numbers TL1TR001437, T32GM080202, T32EY014537, P30EY001931, and R01EY024969. This investigation was conducted in a facility constructed with support from Research Facilities Improvement Program, Grant Number C06RR016511, from the National Center for Research Resources, National Institutes of Health. The content is solely the responsibility of the authors and does not necessarily represent the official views of the National Institutes of Health. This work was also supported by Vision for Tomorrow and the Thomas M. Aaberg, Sr., Retina Research Fund.

---

**Supplemental Figure S1:**
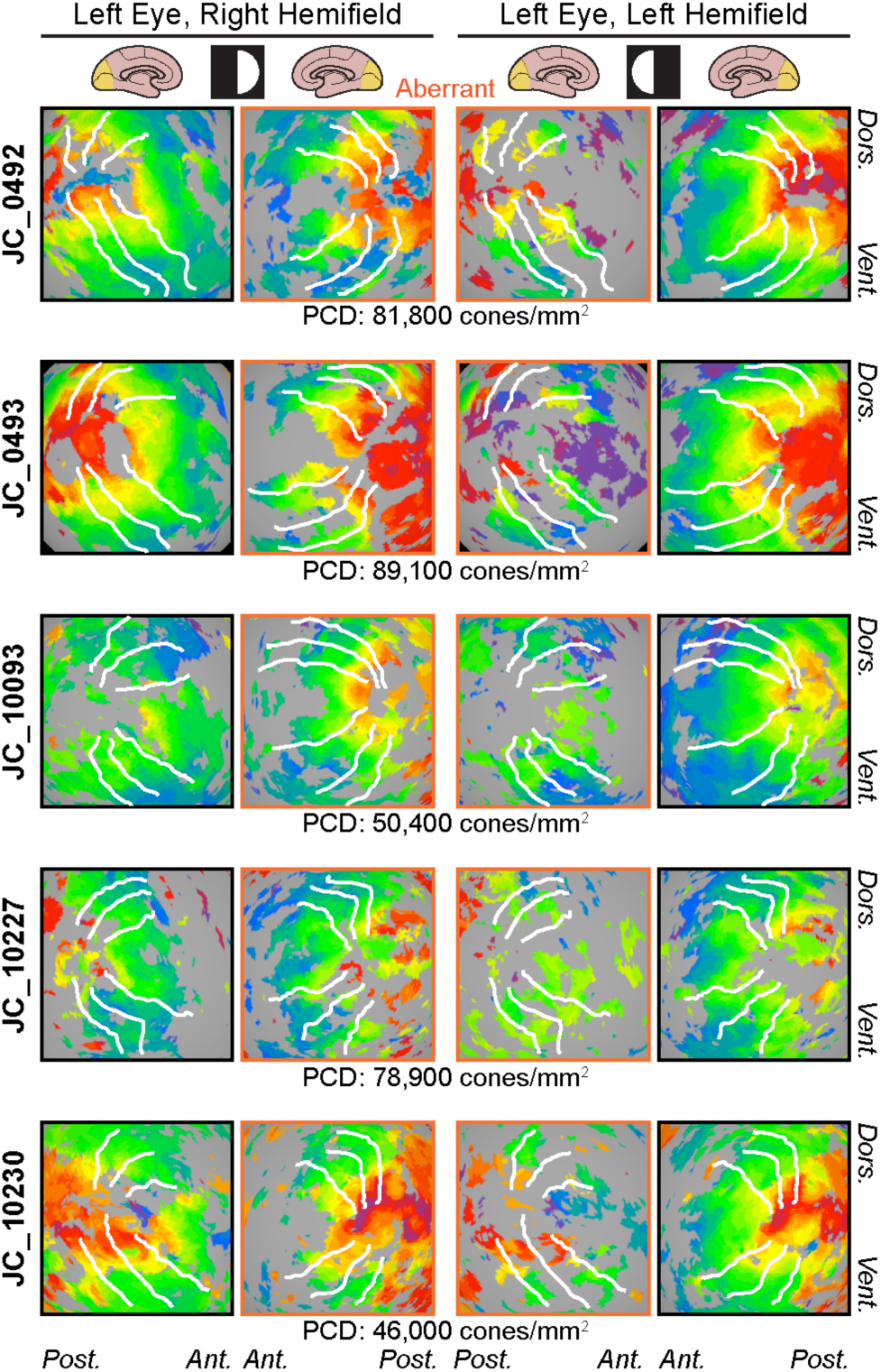
Retinotopic maps of visual field eccentricity for left eye viewing condition in subjects with albinism. Visual stimulus and color coding as shown in Figure 4A. Retinotopy patterns outlined in orange are aberrant ipsilateral hemifield representations. Peak cone densities for each subject with albinism indicated below each row. Maps are displayed on spherically-inflated cortical surface models. Visual field stimuli are indicated by white circle or semicircle symbols at head of respective columns. White lines mark dorsal and ventral boundaries of V1/2/3 based on polar angle data (cf. Figure 1). PCD = peak cone density; Dors. = dorsal; Vent. = ventral; Ant. = anterior; Post. = posterior.

**Supplemental Figure S2:**
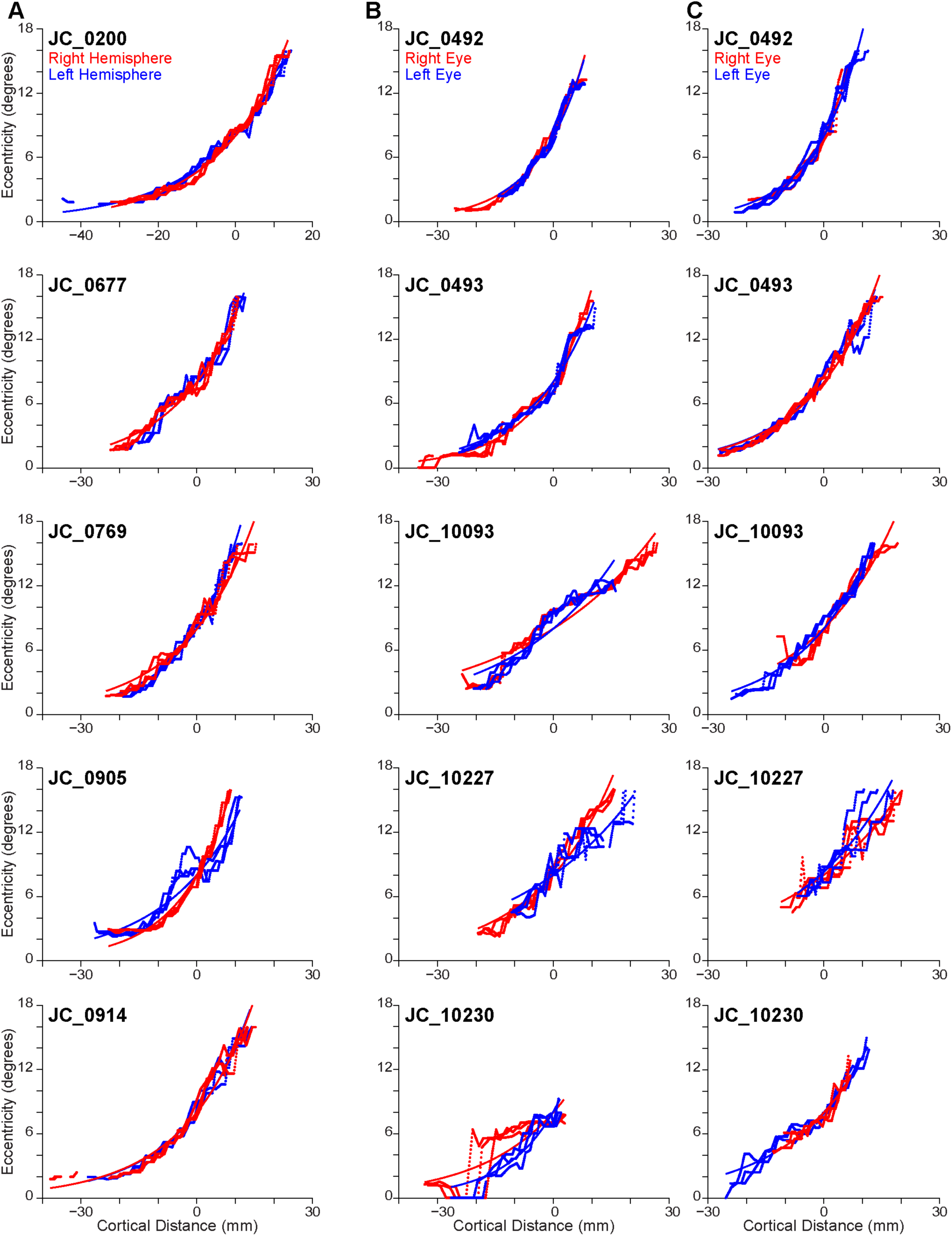
Empirical cortical mapping data and fitted functions for all subjects. (A) Controls: right hemisphere (red) and left hemisphere (blue). (B) Albinism: left hemisphere, right hemifield stimulus. (C) Albinism: right hemisphere, left hemifield stimulus. Viewing conditions in B, C: right eye (red), left eye (blue).

## Notes

CONFLICTS OF INTEREST: The authors declare no competing financial interests.

